# Magnetoencephalography Without a Shielded Room

**DOI:** 10.64898/2026.08.06.743270

**Authors:** Yulia Bezsudnova, Nicholas A. Alexander, Stephanie J. Mellor, Sergey Mitryukovskiy, Rudy Romain, Agustin Palacios-Laloy, Gareth R. Barnes, Martina F. Callaghan, Tim M. Tierney

## Abstract

Magnetoencephalography (MEG) offers non-invasive neuroimaging with high temporal and spatial precision - but its adoption is hampered by the prohibitive cost and infrastructure burden of a magnetically shielded room. We have overcome that burden and present a lightweight, low-cost, multichannel magnetoencephalography system that can image brain activity without needing a magnetically shielded room. The multichannel nature of the system facilitates not just detection but also localization of brain signals that are over 300 million times smaller than environmental interference, without requiring passive shielding. Our system weighs less than 75kg, more than 100 times lighter than a typical shielded room. This is made possible through low-cost active shielding and software-based spatial filtering. We also show that the signal to noise ratio of our in-vivo recordings is comparable to what can be obtained from a conventional cryogenically-cooled MEG system sited within a shielded room. This demonstration is a crucial step towards democratizing magnetoencephalography and making it a globally accessible neuroimaging technology for healthcare and discovery research.

## INTRODUCTION

Magnetoencephalography (MEG) non-invasively measures the magnetic fields generated by brain activity^1^. However, these magnetic fields are more than 300 million times smaller than the Earth’s magnetic field. This necessitates incredibly sensitive sensors to detect the neuronal signals over and above the much larger electromagnetic. interference that pervades the environment. Conventionally, this has been achieved through the use of cryogenically-cooled superconducting quantum interference devices (SQUIDs) operating within a magnetically shielded room^2^. The financial cost and the need to site the multi-tonne shielded room, the MEG system itself (typically >500 kg), and potentially an on-site helium recycler^3^ has limited the widespread adoption of MEG, despite its proven clinical benefit - for example, in predicting surgical outcomes for people with epilepsy^4^.

Optically pumped magnetometers (OPMs)^5–8^, are sensors that have recently been proposed as an alternative to conventional SQUID-based MEG. Their lightweight, cryogen-free design simplifies siting requirements, and means that they can be placed much closer to the brain, boosting signal amplitude^9^. OPMs can additionally be combined with active shielding, which measures the ambient magnetic field using reference sensors and subsequently applies counteracting fields that cancel any interference in real time. Active shielding has benefitted OPM recordings greatly by enabling recordings of freely moving participants^10^, reduced OPM sensor nonlinearities^11^, improved precision of source localization^12^ and deployment OPMs in challenging clinical environments^13^. Furthermore, active shielding can also reduce demands on the passive shielding^11,14–16^, for example by allowing lightweight rooms^17^ to be constructed with fewer layers of mu metal^18^.

Despite these substantial technological advances, the barrier to entry remains the need for a heavy and expensive magnetically shielded room. One approach has been to improve accessibility by making the shielded rooms transportable, albeit while still weighing >8000kg^19^. More desirable still is to make OPM recordings without any shielded room. Attempts to achieve this have delivered incredibly promising results, but have been restricted to single-channel systems with sensitive axes constrained by a single, fixed bias field^20,21^ or low baseline (2 cm) 1^st^-order gradiometers. Although single channel systems can be of great use in brain computer interface^22^ or cardiac imaging^23^ applications, they cannot achieve the spatial localization and whole brain coverage^24–26^ that are critical for both clinical applications and discovery research^1,4^. With regards to low baseline gradiometers, they can offer impressive interference mitigation^27^ but such baselines cause substantial signal loss (∼50%)^28^ and remain vulnerable to any interference that manifests as a large spatial gradient, e.g., as caused by proximity to lifts in an office, lab or hospital environments. Indeed, early work on the optimization of interference mitigation for cryogenic systems suggest that first order gradiometers are insufficient to minimize interference in complex environments and that higher order cancellation is required^29^.

To achieve this higher order level of cancellation without sacrificing signal one can leverage the intrinsic spatial orthogonality of vector magnetometers signals to environmental interference^25,30–32^. There are now a wide variety of post-hoc methods that can remove such high order interference from OPM recordings via software tools (henceforth referred to as “software shielding”)^32–39^. These tools have played an important role in harmonizing results across sites^13^, optimizing source localization^40^ and demonstrating equivalence between OPM and cryogenic MEG systems^41^. Indeed, the flexible combination of active, passive and software shielding has even been used to create person sized shields^42^ that facilitate more portable multichannel recordings. However, when it comes to developing a cost-effective, lightweight and more accessible MEG system, the optimal balance between active, passive and software shielding still remains unclear.

Our previous work has focused on defining the limits of what software shielding can contribute to interference suppression. We have shown that there is a deterministic relationship, regardless of array design, between software shielding (specifically orthogonal projections) and sensor linearity^34^. This relationship can be used to define how well an active shielding system needs to perform in order for passive shielding to be replaced with much lower cost active and software shielding. Based on this, we hypothesize that noise floors commensurate with MEG (10s of 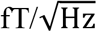) require OPM sensors with linearity errors on the order of 0.1%. In addition, we hypothesize that dynamic ranges of approximately +/-100nT will suffice to compensate for any residual static magnetic fields and their gradients caused by imperfect coil design. This level of performance is crucial for whole head coverage.

Here we show that these criteria can be met by current, commercially available technology to deliver a lightweight, low cost, multi-channel MEG system capable of operating without a shielded room. We successfully spatially localize MEG responses *in vivo* and demonstrate that the SNR of these recordings is comparable to that of conventional SQUID-based MEG that does leverage a shielded room.

## RESULTS

### System Overview and Static Shielding

The system (**Fig. 1A**) has been assembled in an unmodified office space at University College London. This inner city, high interference environment poses a significant challenge such that successful recordings here would provide high confidence for translation to other clinical and research environments. The system consists of three main components: a set of electromagnetic coils that cancel the Earth’s magnetic field, a feedback system that continuously updates the coil currents, and OPM sensors that measure the neuromagnetic field of interest. The coils are split into two sets. The “static coil set” cancels the temporally static components of the Earth’s magnetic field (i.e. provides static shielding). The “dynamic coil set” provides real-time cancelation of the fluctuations (e.g., due to nearby lifts) of the field (i.e. dynamic shielding). We note here that our environment necessitates real-time control of both the field terms and gradient terms, highlighting the need for higher order magnetic field cancellation than first order gradiometers can provide^29^. We use a 48 channel, MAG4Health commercial OPM system^43^ for measurement of the neuromagnetic field. The static shielding (**Fig. 1B**) reduces the average field (and variation across channels) from 55,000 nT (range: 1600nT) to 64 nT (range: 70nT). These measurements lie comfortably within the OPM sensors’ +/-200 nT operational range, enabling whole head coverage.

**Figure 1.**
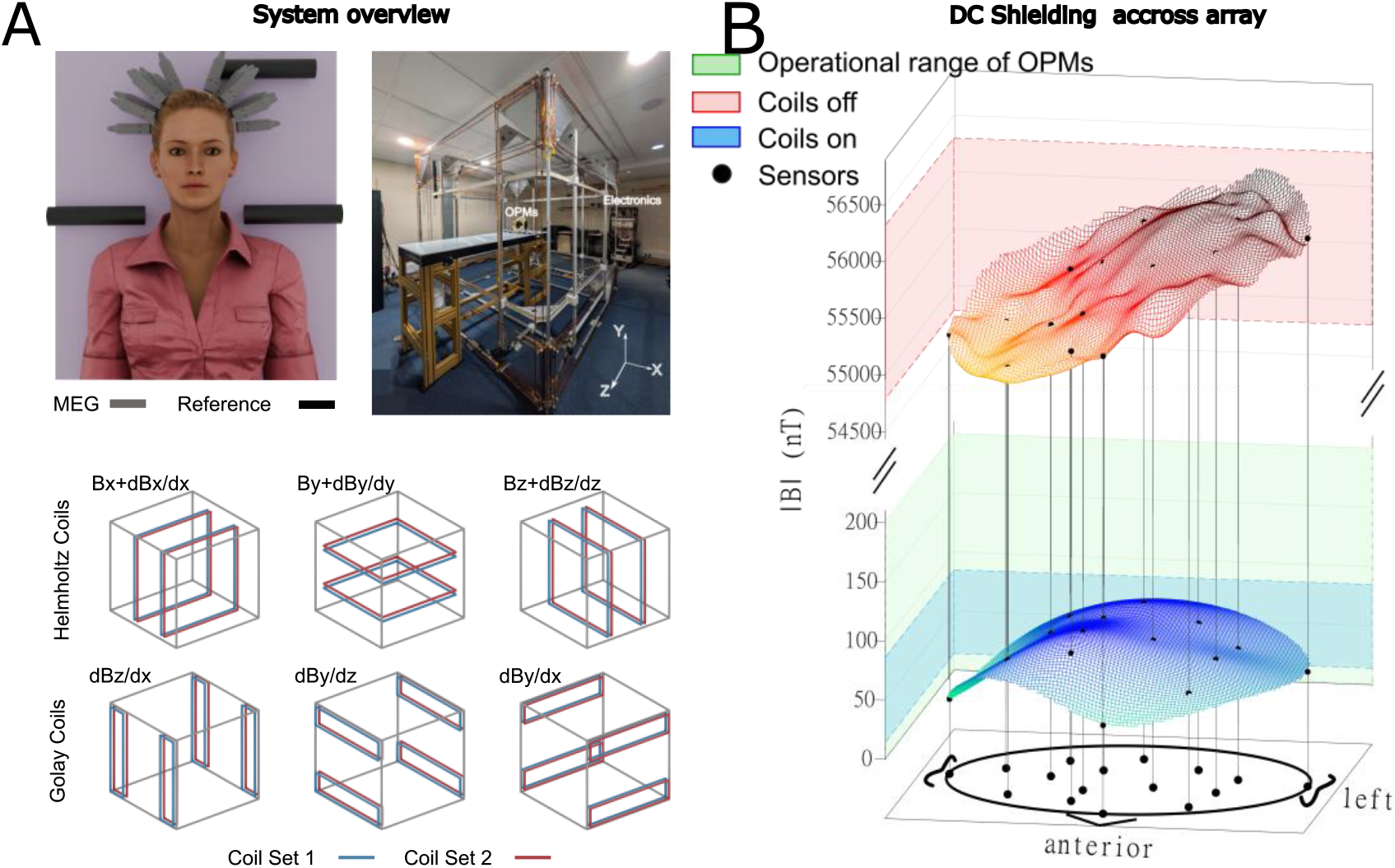
System overview and static shielding. In A) the MEG sensors covering somatosensory cortex are shown on a computer generated avatar. In addition, the reference sensors driving the schematised Helmholtz and Golay coils are also shown. A photograph of the actual setup is also shown. In B) the field norms are shown on the vertical axis when the coils are turned off (red shading) and turned on (blue shading). The operational range of the OPMs is shown in green. The other two axes show how the field norms change in space. Black dots indicate sensor locations across a head sized volume.

### Dynamic Shielding

Once the ambient magnetic field is within the OPM’s operational range, it must be maintained within that range to facilitate MEG recordings. To achieve this we use dynamic shielding^11,15^. As shown in **Fig. 2A**, without dynamic shielding, the field shows peak to peak changes on the order of 1,000 nT (dominated by interference from nearby lifts), well outside the 200 nT operational range of the OPMs. With dynamic shielding and software-based spatial filtering^38^ the standard deviation of the magnetic field is reduced from 258 nT to 0.07 nT. In **Fig. 2B**, the broadband (1 – 80 Hz) shielding factor is greater than 1000 and brings the system from an interference-limited noise floor of 50,000–100,000 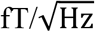to within the manufacturer’s noise floor specification of 45 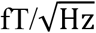 (black dashed line **Fig. 2B**). **Fig. 2C** shows the recovery of a phantom signal with magnitude <400 fT (comparable to a realistic neuronal field) in an environment changing by ∼1 µT over time (**Fig. 2A**), a signal-to-noise ratio (SNR) improvement in excess of 2,000,000. This vast improvement in SNR is also facilitated by the OPM sensor having the requisite low gain error (0.11%, **Fig. 2D**) and thus maximising interference mitigation for spatially complex interference^34^.

**Figure 2.**
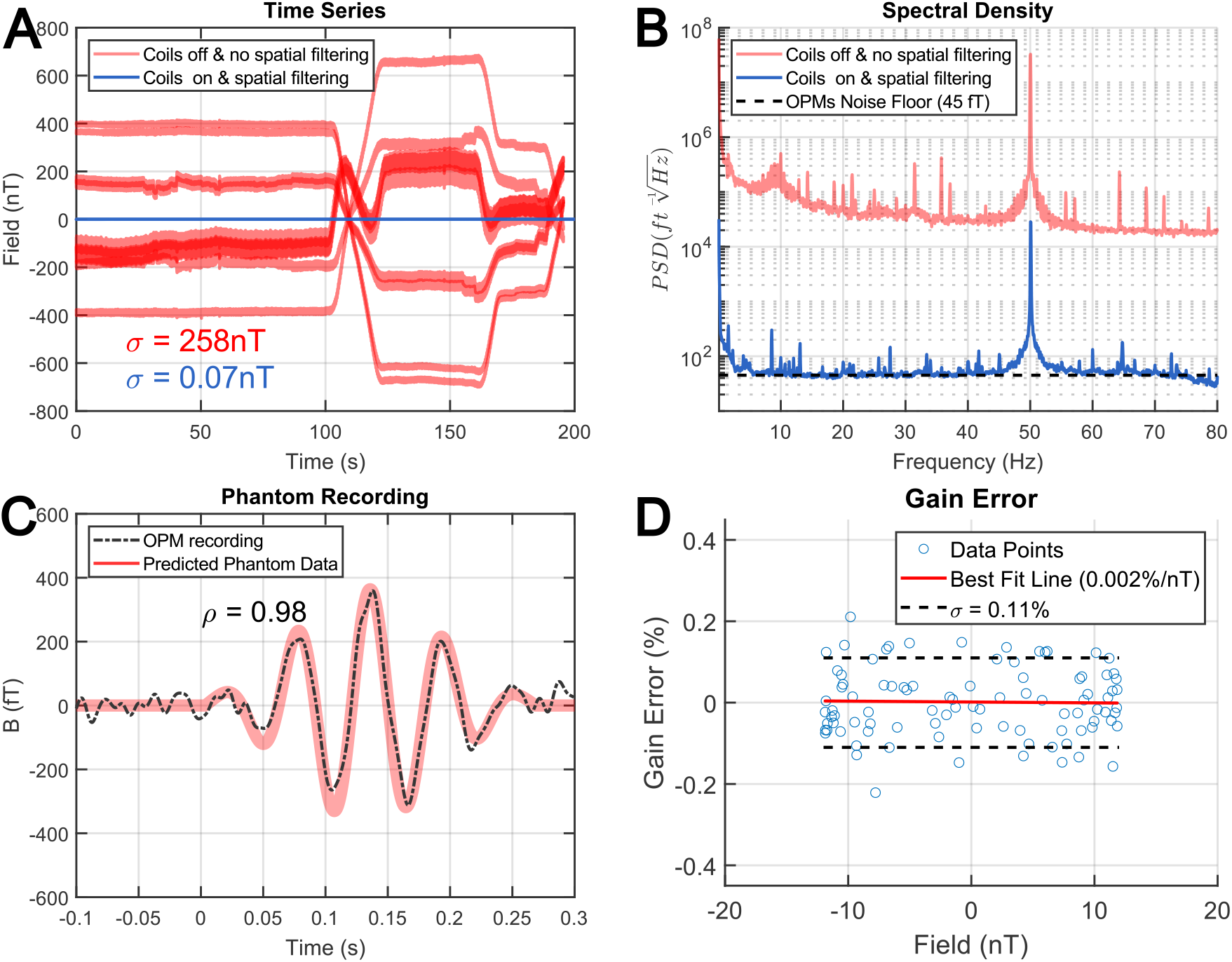
Dynamic shielding and phantom recording. In A) the magnetic field time series (reference sensor) is displayed in red when only the static field cancellation is applied. In blue, OPM recordings are shown when the dynamic cancellation and software-based spatial filtering is applied. In B) the root mean square spectrum is shown as a function of frequency. The dashed black line indicates the OPM noise floor. Panel C) shows predicted phantom data (in red). The observed data is shown via a dashed black line. The correlation, ρ, is indicated in text. In D) the gain error of the OPM system (Y-axis) is shown as a function of field(x-axis) The standard deviation of the gain error is indicated by black dashed lines. The line of best fit relating field to gain errors is shown in red.

### In-Vivo demonstration

As a consequence of minimising both the static and dynamic components of the environment’s magnetic field we now demonstrate our key result: measurement and localisation of neuronal activity without a shielded room. We elicited a somatosensory evoked field from a healthy volunteer by stimulating the left median nerve (∼2Hz stimulation and 2000 trials (2 runs of 1000 trials) – **Fig 3.A**). The OPM array consisted of 16 triaxial sensors (48 channels) positioned over the right (30 channels) and left (18 channels) side of the head to cover the expected field topography of the task (**Fig. 3B**). Our OPM system detected a response at the expected latency of 20ms post stimulation (blue shaded region of **Fig. 3C**) and showed high inter-run correlation (0.93) and high correlation (0.94) with somatosensory lead fields (**Fig. 3D**). The response localised to the expected source location (**Fig. 3E**) of primary somatosensory cortex (MNI coordinate: 35, -26, 46). While all trials were analysed in Fig. 3, we note that the M20 response was detected with statistical significance, controlling for all channels and timepoints (FWE p< 0.05), in 220 trials.

**Figure 3.**
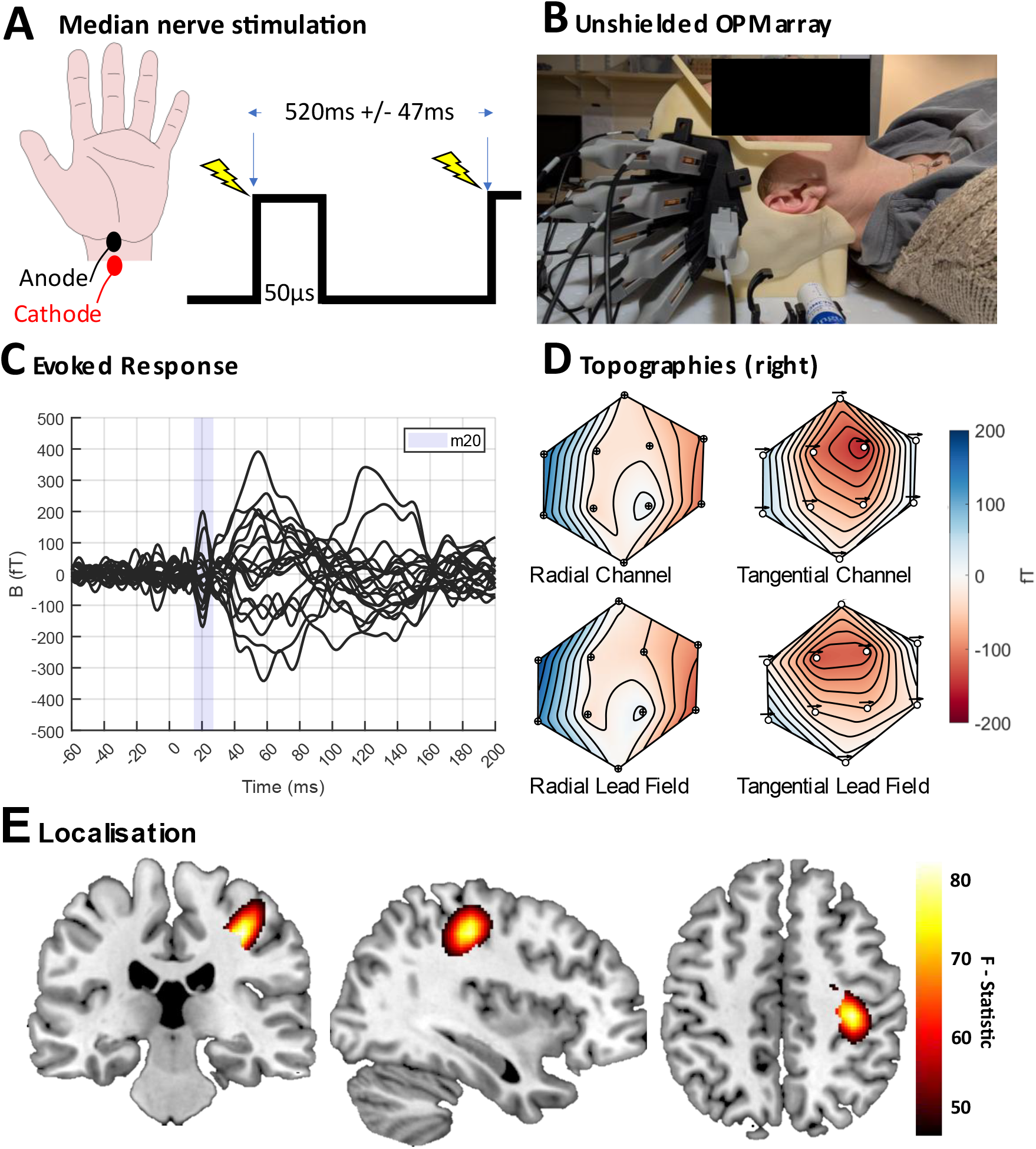
MEG without a shielded room. **A)** shows median nerve stimulation protocol. In **B)** the positioning of sensors over the right sensorimotor cortex is shown (note a further 6 sensors are positioned over left somatosensory cortex).The photo is an obscured photo of the first author. In **C)** the evoked response is shown (for the channels over the right somatosensory cortex). Time zero is the stimulation onset and the blue shaded region shows the expected time period of the M20 response. In **D)** the measured radial and tangential topographies (top row) and lead fields (bottom row) of the sensors visible in **B)** are shown. The tangential sensing direction (posterior to anterior) is shown with arrows over the sensor position. In **E)** the M20 response is localised to primary somatosensory cortex. The statistical parametric map (F-contrast) is thresholded at half maximum and displayed on an MNI template.

### Reference Measurement within a Shielded Room

In addition to localizing the expected M20 response we also examined how the sensor level SNR of our OPM system compared to that of a SQUID gradiometer system (radial gradiometer with synthetic third order gradiometry applied) using the same paradigm with the same participant. We chose the SQUID sensor with the highest SNR as a comparator to our OPM system. For the OPM system we chose the highest SNR synthetic gradiometer (weighted difference of two OPM channels after spatial filtering is applied). While the OPM system showed higher signal amplitude than the cryogenic system it also showed higher variability (**Fig. 4A)**. As such, we compared both systems in terms of their M20 SNR (**Fig. 4B**). We estimated the variability in the SNR via a bootstrapping procedure and replicated this comparison across all trials. The SNR was compared favourably across both systems despite the absence of passive shielding in the OPM case (OPM SNR = 10.85, CI = 8.84 – 12.76, SQUID SNR = 13.6, CI =11.53 – 15.36). The SNR of the OPM data in the unprocessed case (no software shielding) had 95% intervals that overlap with 0 even after 1000 trials of averaging (Unprocessed OPM SNR = -0.74, CI = -2.6 – 1.16). This demonstrates the crucial nature of software shielding for achieving suitable levels of SNR for MEG recordings in humans outside a shielded room. We note here that due to high levels of gradient interference in our environment the higher order cancellation provided by the spatial filtering is crucial to achieving higher SNR^29^.

**Figure 4.**
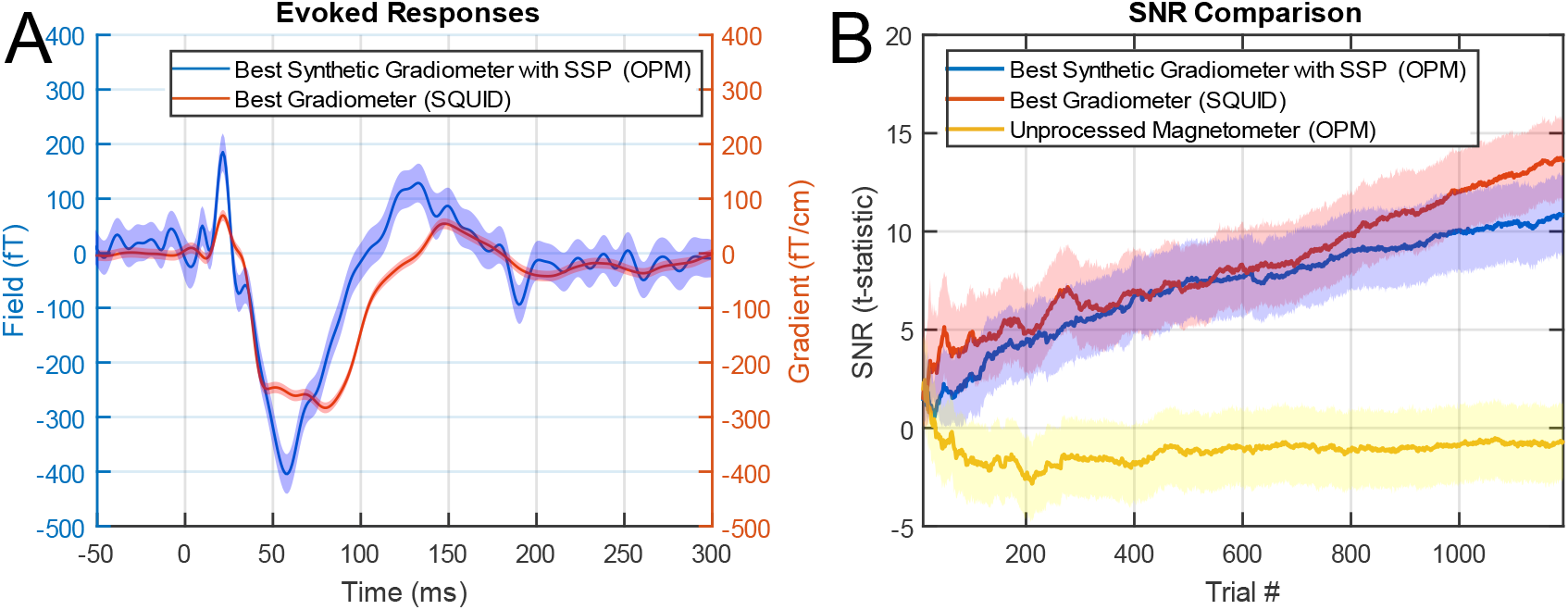
Comparison with a shielded gradiometer system. In **A** the evoked response from the shielded gradiometer with the highest M20 SNR is shown in red. The corresponding highest SNR synthetic gradiometer (weighted difference of two OPM channels after spatial filtering is applied) evoked response from the OPM system is shown in blue. Shaded regions indicate 95% confidence intervals. In **B** the SNR (expressed as a t-statistic) of the M20 response is shown for both systems and for the unprocessed OPM data (processed OPM – blue, SQUID – red, unprocessed OPM data - yellow) as a function of trial number. Bootstrapped estimates for the 95% confidence intervals are shown as shaded regions

## DISCUSSION

We have shown that standard, commercially-available, multi-channel vector OPM systems can be used to both detect and localize neuronal signals without using a magnetically shielded room. We achieve this by combining active shielding with additional software shielding to detect neuronal signals that are ∼300 million times smaller than the background magnetic field. We also note here that the ambient field changes we observe (∼1000nT at low frequency and ∼300 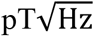 at 10Hz) are more than 20-50 times greater than what has been reported in other studies of OPMs operating without shielded rooms^20,21,23,27^. Despite, the comparatively minuscule size of the brain signal compared to these changes, the SNR of our *in vivo* recording compares favorably with state-of-the-art SQUID-based MEG recordings made within a shielded room. This emphasizes the potential of our approach to democratize access to MEG at a truly global level, without compromising on data quality.

Reducing the passive shielding (and therefore financial and siting) requirements of MEG has been a crucial area of research to enable the propagation of this technology. Cryogenic systems have demonstrated remarkable robustness to magnetic interference but still weigh many hundreds to thousands of kg, limiting their portability^3^. In contrast, OPMs are far more lightweight (a few hundred grams) but as they have no inbuilt mechanism to reject interference, their multichannel implementation, even in portable applications, has typically required a fully shielded room^19^ (weighing 8,000kg). Our active shielding framework eliminates this limitation.

For single channel OPM systems there has been great progress in operating OPMs outside of shielded rooms by using bias fields^20,21^. However, such implementations result in measuring the projection of the brain’s field along that bias field as opposed to an optimal sampling of the neuronal signal. As such, approaches that only measure in one direction suffer from a similar issue faced by radial only magnetometer systems: They struggle to spatially separate brain signal from spatially complex interference or dynamically changing gradients^25,26,35^. Similarly, low baseline gradiometers^27^ without higher order cancellation also suffer from this issue^29^. We address this issue by using vector magnetometers that are highly optimized for separating brain signal from interference^25^. We note that in our environment, due to extensive gradient noise, the SNR of synthetic gradiometry, without higher order spatial filtering results in five times lower SNR than when spatial filtering is used. This spatial separation of brain signal from spatially complex interference is crucial for generalization of this technology to challenging magnetic environments^29^ (anecdotally, our system is surrounded by 3 MRI scanners, 2 lifts, sits above the London underground and has building works taking place above it). This ability of our system to separate large, complex interference from brain signal is fundamental to achieving SNR comparable to the fully shielded system.

While the system proposed here is more than 100 times lighter than existing shielded rooms (70Kg vs 8000Kg) the coils we utilized in the current setup (square Helmhotz and Golay coils) are large in size (2m x 2m x 2m) and would require 1-2 days to be disassembled and reassembled if portability was required. As such our next step would be to take advantage of target field coil design methods^44,45^ to miniaturize the coil system substantially and enable truly portable MEG in any environment. This would not only minimise cost in resource constrained settings but also open up new opportunities in areas such as community based neurological assessment^46^ and sports related brain injury^47^

The success of a truly portable system will be predicated on achieving signal quality sufficient for MEG recordings. As such, our final analysis compared sensor level SNR between our unshielded OPM system and a shielded SQUID gradiometer. While it is incredibly encouraging that the two systems showed similar SNR such comparisons could be done in a multitude of different ways and are incredibly non-trivial to perform. For instance, the experiments here were conducted on separate days and while the stimulation was calibrated each day on the same person, we acknowledge this as a potential confound. Furthermore, the OPM system has at least 3dB of signal suppressed due to the number of channels available (48) and the spatial filtering used^38^. As such, future work will focus on both expanding the system to have more channels while simultaneously optimizing our spatial filtering methods. This would enable more detailed quantitative comparisons between shielded SQUIDs and unshielded OPMs, similar to comparisons of OPM recordings within shielded rooms to SQUIDs that have already ben made^7,41^. The presented analysis therefore serves as a useful reference point for benchmarking the performance of our proposed system as opposed to a definitive technical comparison of the two systems

In summary, we have addressed the challenges of real-time interference mitigation and suppression of the Earth’s magnetic field without the need for passive shielding in the form of a shielded room. In doing so, we demonstrate that lightweight, cryogen free, multichannel MEG systems can be feasibly deployed in real-world environments to both measure and localise brain activity. This work capitalizes on the rapid progress in MEG system development over the past decade. In particular, the transition from bulky cryogenic MEG systems to lightweight, actively shielded^11,14^ OPMs has laid the foundations for MEG without expensive magnetically shielded rooms. By eliminating such key financial and infrastructural barriers, our approach opens MEG to broader use - transforming it from a specialized research tool into a globally accessible technology for healthcare and discovery research.

## MATERIALS AND METHODS

### System overview

The system consists of three main components: a set of coils that cancels the static Earth’s magnetic field, a feedback system that controls the current supplied to these coils and OPM sensors that measure the brain’s neuromagnetic field. The coil configuration includes six Helmholtz coils, which cancel the spatially uniform components of the ambient magnetic field as well as its longitudinal gradients: 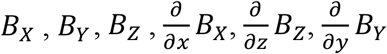. Among these 6 coils, three coils counter the static (average in time) components of the field, while the other three compensate dynamic fluctuations in these fields and gradients over time. The Helmholtz coils (Ferronato BHC200-3-B) each measure 2m × 2m and consist of 16 turns. Their field and gradient efficiencies are 14.4 μT A^-1^ and 13.0 μT A^-1^m^-1^, respectively.

An additional set of six Golay^11^ coils is used to control the transverse gradients: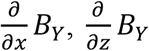, and 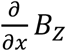. Three of these coils counter the static components of the transverse magnetic field gradients, while the other three compensate for their dynamic fluctuations. Each Golay coil section comprises 10 hand-wound loops of enameled copper wire (1.18mm diameter), with dimensions of 0.34 m × 2.04 m, resulting in a gradient efficiency of 1.8 μT A^-1^m^-1^.

The coils are controlled via a feedback system of 8 power supplies (for static field control) and 8 analogue PID (Proportional, Integral, Derivative) controllers (for dynamic field control) that minimise the field on 3 reference vector fluxgate magnetometers.

The OPM sensor array used to measure brain signals in the current study is a commercially available triaxial 16 sensor (48 channel) helium-based OPM system, built by MAG4Health^7,43,48^. This OPM system achieves a closed loop dynamic range of ±200 nT with gain error = 0.11%. The sensitivity is at most 45 fT/√Hz on two out of three axes (one radial to the head and one tangential) and <300 fT/√Hz for the third axis. The high dynamic range (+/-200 nT) and stable linearity makes this system suitable for recordings in environments with large environmental interference.

### Static shielding

Three tri-axial Bartington MAG-03 -MC fluxgate sensors are used to measure the background magnetic field. They have a noise floor of approximately 7pTrms/√Hz and measuring ranges of ±70μT. They are positioned in a triangular formation (30cm separation - **Fig. 1A**) around the OPM system so as to measure the three static (in time) uniform field components, three independent transverse gradients and two independent longitudinal gradients. The analogue outputs from these sensors are digitised using a 16-bit National instruments ADC (NI-9205) and read into a custom, real-time LabView data acquisition program. Once the field and gradient terms are measured, the current required to counteract and minimise these terms is determined automatically (typically under 30s) using a gradient descent algorithm on the sum of squares of the fluxgate outputs. This procedure is terminated when the field is less than 30nT on each axis (well within the 200nT dynamic range of our OPM sensors). The calculated current is supplied to the coils using three BK Precision 9206B and two Aim-TTi MX100TP power supplies.

To evaluate the static shielding performance of the system (**Fig. 1B**), we compared the temporally invariant component of the magnetic field distribution around the OPM sensors with the shielding coils either switched off or on. To estimate the field with the coils off, we used a field-mapping technique^49^ in which the position of a fluxgate magnetometer was tracked using optical motion capture of passive, retroreflective markers (OptiTrack DUO, NaturalPoint Inc., OR, USA) while the fluxgate sensor was manually scanned across the area of interest. With the coils on, we recorded the outputs directly from the OPMs mounted on the helmet with whole head coverage. As the field is constantly changing in time, we performed these measurements while a feedback system (described in the next section) was used to minimise the dynamic changes in time. In this way, our measurements were only a function of the static components of the field. Note that in the coils off case the field was not sampled by the fluxgate at the position of the OPM and was therefore linearly interpolated in this case.

### Dynamic Shielding

Once the gradient descent step of the static shielding stage is complete, dynamic control is activated to maintain magnetic field stability over time. This is achieved by feeding eight of the fluxgate analogue outputs (8 degrees of freedom are required to control the 3 uniform field components and 5 independent first order gradients) into eight analogue PID controllers (SRS SIM960). PID terms are tuned manually to minimise the standard deviation of the signal in time. These PID controllers dynamically supply currents of up to 100 mA in real-time. This allows for dynamic uniform field, longitudinal gradient, and transverse gradient control of +/-1.4 μT, +/-1.3 μT m^-1^ and 0.18 μT m^-1^ respectively. Should the output of the PIDs approach saturation (monitored in real-time via a custom Labview program), we automatically adjust the current on the static shielding coils using a digital PID controller to update the BK Precision 9206B and Aim-TTi MX100TP power supplies. This forms an effective ‘cascaded’ or ‘chained’ PID control system and ensures continual operation of the system even during periods of excessively large dynamic environmental disturbance (>1.4 μT).

As the dynamic shielding is driven by reference sensors with noise floors of ∼7pTrms/√Hz, this random electronic noise creates spatially structured magnetic interference at the locations of the OPMs. For all OPM recordings we use Signal Space Projection^38^ (SSP) to construct a spatial basis set of interference vectors from one minute of empty room OPM recordings, taken prior to each experimental recording. This spatial basis set is derived using all 48 channels and a weighted least squares approach is used to regress the interference from the data. Note that as the hardware component of the dynamic shielding is limited by the fluxgate noise floor, we report the combined shielding effect of both software and coils in **Fig. 2A & 2B**.

Two separate recordings were made to evaluate the performance of the dynamic shielding. In the first recording the static shielding was applied but not the dynamic shielding. Data were recorded from the 3 reference triaxial fluxgates (because the environmental magnetic field changes are larger than the operating range of the OPM system). In the second recording, data were acquired from the 48 channel OPM system with the dynamic shielding turned on. We also applied the SSP projection to the OPM recordings to show the combined impact of the coils and spatial filtering (**Fig. 2A & B**).

### Phantom Recording

Phantom recordings were performed by placing a helmet containing 16 evenly distributed (∼8cm nearest neighbour distance) OPM sensors (48 channels) at the centre of our coils. To create a brain-like signal, we used a small coil (3 mm radius and 10 turns), displaced 3.8 cm from the cell of the nearest OPM sensor. A ∼1 μA (∼ 30 nAm^2^ moment) oscillating current was applied to the coil to generate a magnetic field of approximately 400 fT at the location of the nearest OPM sensor cell. The stimulus waveform was generated using a microcontroller (Arduino Uno R4). It consisted of a 17 Hz sinusoidal signal multiplied by a Hamming window with a duration of 270 ms. The inter-stimulus interval (ISI) was 250 ms ± 47 ms. We recorded one minute of data without the phantom signal, followed by ten minutes with the phantom signal active.

To analyse the phantom recordings, the data were first band-pass filtered between 1 and 80 Hz, with an additional notch filter applied between 47 and 53 Hz as well as 97 and 103 Hz to suppress line noise and its first harmonic. SSP was applied to minimise environmental interference, using interference components derived from the minute long recording without the phantom signal active. The data were then epoched from −100 ms to 300 ms relative to stimulus onset, baseline-corrected using the pre-stimulus interval (−100 ms to 0 ms), and subsequently averaged (**Fig 2D**).

### Gain assessment

To assess the stability of sensor gain, we used the same phantom as described in the previous section to generate an approximately 15 nT reference sinusoid at 17 Hz and recorded OPM data for 100 s. We then broke this OPM data into 500 ms overlapping blocks and measured the Fourier amplitude of this 17 Hz sinusoid in each block. The gain error was calculated as the ratio of the standard deviation in the reference signal’s amplitude relative to its mean, over time.

### Participants

We recruited one healthy, right-handed participant to take part in this experiment (1 Male; age 59 years), who provided written, informed consent. The study procedures were approved by the University College London Research Ethics Committee and experiments were conducted in accordance with the Declaration of Helsinki.

### Paradigm

The somatosensory paradigm involved electrical stimulation of the participant’s left median nerve. 50 µs-duration current pulses were applied to electrodes placed on the participant’s wrist (**Fig. 3A**) using a Digitimer DS7A constant current stimulator. The current amplitude was increased until a visible thumb movement was observed. Pulses were delivered at an inter-stimulus interval of 520 ms with a uniform jitter of +/-47 ms. Each of the 2 experimental runs lasted 10 minutes (around 1000 trials). Before the experimental run, a 2-minute empty-room recording was acquired without the participant present. The same paradigm was performed in the shielded room, on a separate day, with the same participant using a CTF 275-channel whole-head CTF MEG system (MISL, Port Coquitlam, Canada) operating in third-order synthetic gradiometer configuration

### Sensor layout

The participant was lying on the bed as shown in **Fig. 1A**, with the head positioned at the centre of the system. The helmet used for the experiment was custom-moulded to match the participant’s head shape using their structural MRI^50^ (**Fig. 3B**). Two hexagonal arrays of slots for up to 20 OPMs were positioned over the participant’s right and left somatosensory cortices. The right-hand slot was populated with 10 OPMs (as the response is lateralised to the right somatosensory cortex) while the left-hand slot was populated with the remaining 6 sensors. The target locations and orientations were estimated from previous SQUID-MEG dipole fits obtained from the same participant.

### Sensor level analysis

To analyse the data, a 1–80 Hz band-pass filter was first applied to both the in-vivo and empty room recording (i.e., with and without the participant present). Notch filters at 47–53 Hz and 97–103 Hz were then applied to suppress line noise and its first harmonic. SSP based on the empty room recording was used to suppress environmental interference from the *in vivo* measurement. The choice of number of interference vectors to use in the SSP step is a trade-off between signal suppression (due to using more degrees of freedom) and improving SNR via increased interference mitigation. Importantly, for *in vivo* and phantom recordings this trade-off can be examined *a priori* by applying the projection learned from the empty room data to the leadfields. We therefore set the number of vectors as the number that would produce <3dB of signal suppression (similar to temporal filtering).

Once SSP was applied, the data were then epoched from −100 ms to 300 ms relative to stimulus onset, baseline-corrected using the pre-stimulus interval (−100 to 0 ms), and averaged. The time series can be seen in **Fig. 3C** while the topographies for the right-hand radial axis and the tangential axis with the lower noise floor are shown in **Fig. 3D**. To assess reproducibility the correlation of the time series between run 1 and run 2 was calculated. Furthermore, the SNR of the response (t-statistic) was examined as a function of trial number, with and without the interference correction applied. Bonferroni correction (correcting for number of channels and timepoints) was used to determine the minimum number of trials required to detect the M20 response.

The SQUID data was filtered using the same frequency filters as the OPM data and epoched to the same time windows. The SQUID sensor with the highest M20 SNR response was selected as the comparator for the OPM data. To aid the comparison of the best radial gradiometer(with synthetic 3^rd^ order gradiometry applied) SQUID data with the OPM data, the synthetic gradiometer (weighted difference of two OPM channels, after SSP) that provide the best SNR for the OPM data was used as a comparator for the SQUID data (**Fig. 4**). The purpose of this analysis is not to provide a detailed tehchnical comparison of the two systems, as this is beyond the scope of this study, but to provide a reference SNR level for the OPM data in order to demonstrate viability of the proposed approach.

### Source level analysis

A dipole fit was performed using SPM^51^ and the DAiSS toolbox (https://github.com/spm/spm/toolbox/DAiSS). The forward model was the single shell model^52^. The dipole fit was performed exhaustively for each voxel on 5×5×5mm^3^ grid in the participant’s MRI space. The variance explained by the dipole at each voxel was converted to the corresponding F-statistic to create a statistical parametric map. For display purposes this map was registered to, and displayed on, the MNI template brain. The image is thresholded at half maximum (**Fig. 3E**).

### Software

All data were analysed with a mixture of custom MATLAB code and the Statistical Parametric Mapping (SPM)^51^ package (https://github.com/spm/spm)

## Funding

Epilepsy Research Institute Fellowship [FY2101].

Wellcome Discovery Research Platform for Naturalistic Neuroimaging [226793/Z/22/Z].

National Brain Appeal [NBA-IF 4]

## Author contributions

Conceptualization: M.F.C., G.R.B., T.M.T., Methodology: Y.B., S.J.M., R.R., S.M., A.PL., T.M.T. Visualization: NA, YB, TMT. Writing: All

## Competing interests

S.M, R.R, A.PL work for Mag4Health, a company which sells OPM sensors. They contributed to the development of the OPM system used here, advised on its operation and helped build the participant support. No other authors declare competing interests.

## Data, code, and materials availability

Code used to analyze the data is freely available in the SPM (https://github.com/spm/spm) software package. Data will be made available upon publication.

